# Fluid flow overcomes antimicrobial resistance by boosting delivery

**DOI:** 10.1101/2024.05.08.591722

**Authors:** Alexander M. Shuppara, Gilberto C. Padron, Anuradha Sharma, Zil Modi, Matthias D. Koch, Joseph E. Sanfilippo

## Abstract

Antimicrobial resistance is an emerging global threat to humanity. As resistance outpaces development, new perspectives are required. For decades, scientists have prioritized chemical optimization, while largely ignoring the physical process of delivery. Here, we used biophysical simulations and microfluidic experiments to explore how fluid flow delivers antimicrobials into communities of the highly resistant pathogen *Pseudomonas aeruginosa*. We discover that increasing flow overcomes bacterial resistance towards three chemically distinct antimicrobials: hydrogen peroxide, gentamicin, and carbenicillin. Without flow, resistant *P. aeruginosa* cells generate local zones of depletion by neutralizing all three antimicrobials through degradation or chemical modification. As flow increases, delivery overwhelms neutralization, allowing antimicrobials to regain effectiveness against resistant bacteria. Additionally, we discover that cells on the edge of a community shield internal cells, and cell-cell shielding is abolished in higher flow regimes. Collectively, our quantitative experiments reveal the unexpected result that physical flow and chemical dosage are equally important to antimicrobial effectiveness. Thus, our results should inspire the incorporation of flow into the discovery, development, and implementation of antimicrobials, and could represent a new strategy to combat antimicrobial resistance.

## Main Text

Antimicrobial resistance is a severe threat to human health^1^. Fundamentally, there are three problems: our supply of effective antimicrobials is dwindling^1^, the development of new antimicrobials lacks profitability^2^, and new antimicrobials will likely succumb to resistance. As obvious solutions to antimicrobial resistance are not emerging, new ideas are required. One potential idea is to focus more on how antimicrobials are delivered to target host microenvironment^3,4^. While antimicrobial delivery is typically dependent on host fluid flow, the effect of flow on antimicrobial delivery is largely unknown.

Mechanical features of the environment determine bacterial behavior and survival^5–9^. In recent years, surface association and fluid flow have been shown to impact a wide range of bacterial processes, such as adhesion^10–13^, motility^14–16^, signaling^17–23^, and virulence^24–28^. While researchers have focused primarily on mechanical forces^29^, research has demonstrated that flow-based transport has large impacts on chemical microenvironments^17,18,30^. For example, flow can shape the local concentrations of metabolites^31,32^, signaling molecules^30^, and chemical stressors^17,33,34^. However, the impact of flow on local concentrations of antimicrobials has not been emphasized.

As flow drives antimicrobial delivery and shapes chemical microenvironments, we reason that studying how flow impacts antimicrobial effectiveness should yield useful discoveries. Specifically, we predict that flow will rapidly replenish antimicrobials and overwhelm bacterial resistance mechanisms that rely on chemical modification or destruction. Here, we implement a strategy that combines biophysical simulations and microfluidic experiments to study how antimicrobials inhibit the highly resistant pathogen *Pseudomonas aeruginosa*. Our work connects the growing problem of antimicrobial resistance to the emerging field of mechano-microbiology and reveals how flow can overcome resistance by boosting delivery.

### Flow enhances H_2_O_2_ delivery into resistant bacterial populations

To learn how flow impacts antimicrobial delivery, we examined how the host-generated antimicrobial hydrogen peroxide (H_2_O_2_)^35^ is delivered into populations of *P. aeruginosa*. *P. aeruginosa* is resistant to H_2_O_2_ through multiple catalases and peroxidases^17,36^. Using custom-fabricated one meter long microfluidic channels (Figure S1), we aimed to gain spatial understanding of H_2_O_2_ delivery (Figure 1A). To guide our experimental design, we first simulated the effect of flow on H_2_O_2_ delivery. Our simulations included three key parameters: flow, H_2_O_2_ diffusion, and H_2_O_2_ removal by cells. By holding diffusion and removal constant, we learned how increasing flow promotes delivery deeper in the simulated resistant population (Figure 1B). These simulations support our model that flow enhances H_2_O_2_ delivery and generate testable hypotheses regarding the interaction between flow and H_2_O_2_ concentration.

**Figure 1:**
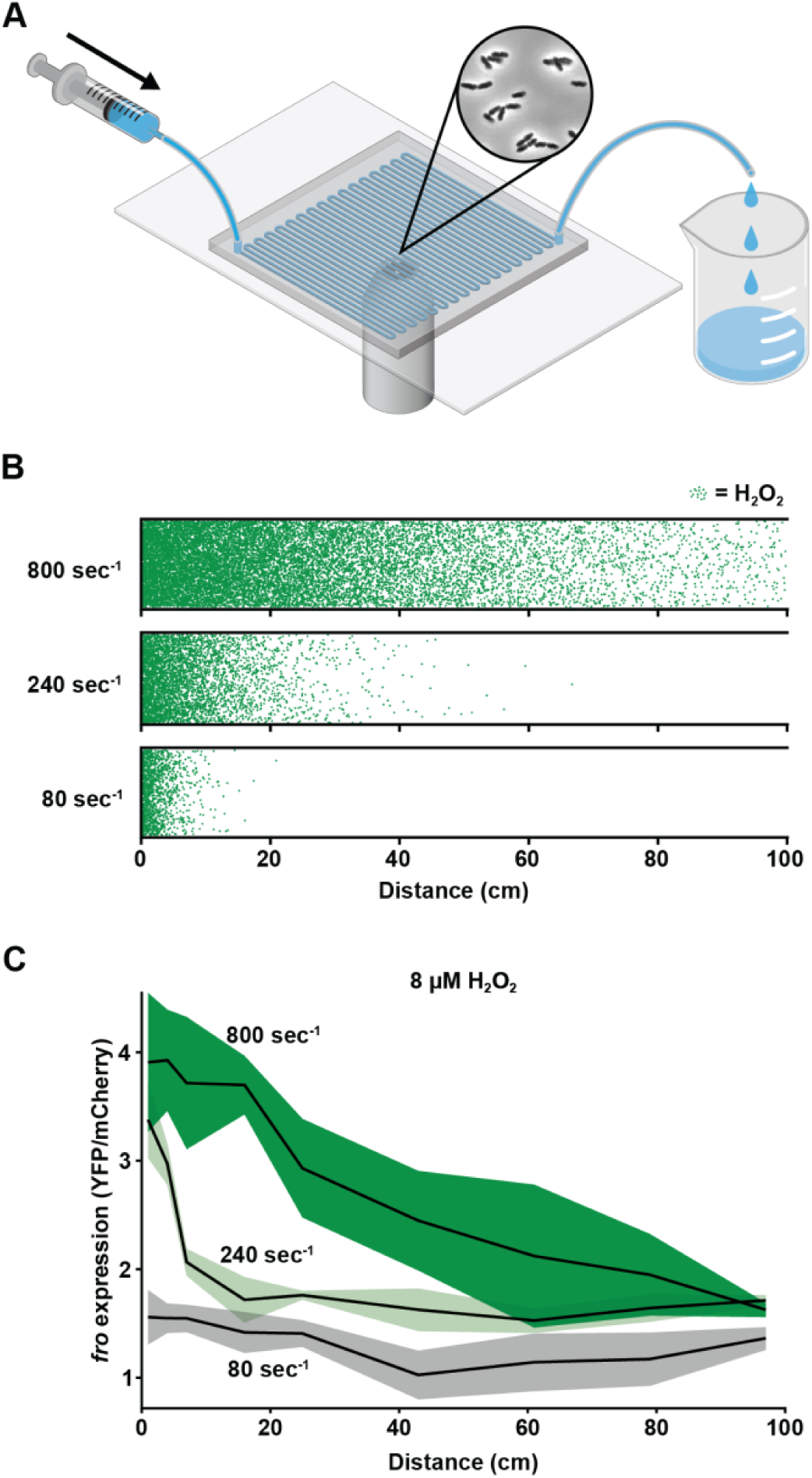
Flow enhances H_2_O_2_ delivery into resistant bacterial populations. **(A)** Illustration of the microfluidic setup that allows for spatial analysis of bacterial populations. Channels are 50 µm tall, 500 µm wide, and 100 cm long. Bacteria adhere naturally to the glass surface. Flow was generated with a syringe pump and images were taken at various distances across the channel. **(B)** Simulation of H_2_O_2_ molecules delivered at different flow intensities (shown as shear rates) in simulated microfluidic devices. Green dots represent individual H_2_O_2_ molecules that experienced diffusion, flow from left to right, and removal when they hit the bottom of the channel to represent bacterial removal. (**C)** *P. aeruginosa fro* reporter expression (which serves as a H_2_O_2_ biosensor) after 3 hours at different shear rates with LB media containing 8 µM H_2_O_2_ in a microfluidic channel. The *fro* experiments in C closely matched the biophysical simulations in B. Shaded regions show SD of three biological replicates.

To experimentally test how flow impacts H_2_O_2_ delivery, we utilized our H_2_O_2_-sensitive *fro* reporter strain. Previous work established that *fro* reporter activity increases proportionately to H_2_O_2_ concentration and thus the *fro* reporter can function as a H_2_O_2_ biosensor. To improve our tracking of biosensor cells, we used a Δ*pilA* mutant version that is unable to move on surfaces but retains indistinguishable H_2_O_2_ sensitivity^18^ (Figure 1C, S2). We seeded our one meter long microfluidic channel with *fro* reporter cells and subjected those cells to 8 µM H_2_O_2_. For these experiments, we used a syringe pump to deliver H_2_O_2_ at varied shear rates (80, 240, 800 sec^-1^) that are commonly found in the human body^37^. Consistent with our simulations, our experimental results show that increasing shear rate increased *fro* reporter activity and lengthened H_2_O_2_ spatial gradients throughout the channel (Figure 1C, S3). Collectively, our simulations and experiments establish that flow enhances H_2_O_2_ delivery into resistant populations of *P. aeruginosa*.

### Flow overcomes bacterial resistance to H_2_O_2_ and gentamicin

How does flow impact bacterial resistance to H_2_O_2_? As H_2_O_2_ resistance relies on chemical degradation through catalases and peroxidases^38^ (Figure S4), we reasoned that increasing delivery with flow could overcome bacterial resistance. To test our hypothesis, we treated *P. aeruginosa* cells with varying H_2_O_2_ concentrations with and without flow. Without flow, cells were resistant to H_2_O_2_, as concentrations up to 64 µM had no impact on growth (Figure 2A, 2B). With flow, cells became sensitive to H_2_O_2_, as concentrations as low as 8 µM impacted growth and a dose of 16 µM completely blocked growth (Figure 2A, 2B). H_2_O_2_ acts as a bacteriostatic antimicrobial, as flowing in media without H_2_O_2_ can restore growth (Figure S5). These results suggest that H_2_O_2_ in flowing host environments restricts bacteria growth, as host H_2_O_2_ concentrations are typically in the low micromolar range^17,35^. Thus, our results reveal that flow restores sensitivity of the resistant human pathogen *P. aeruginosa* to host-relevant concentrations of H_2_O_2_.

**Figure 2:**
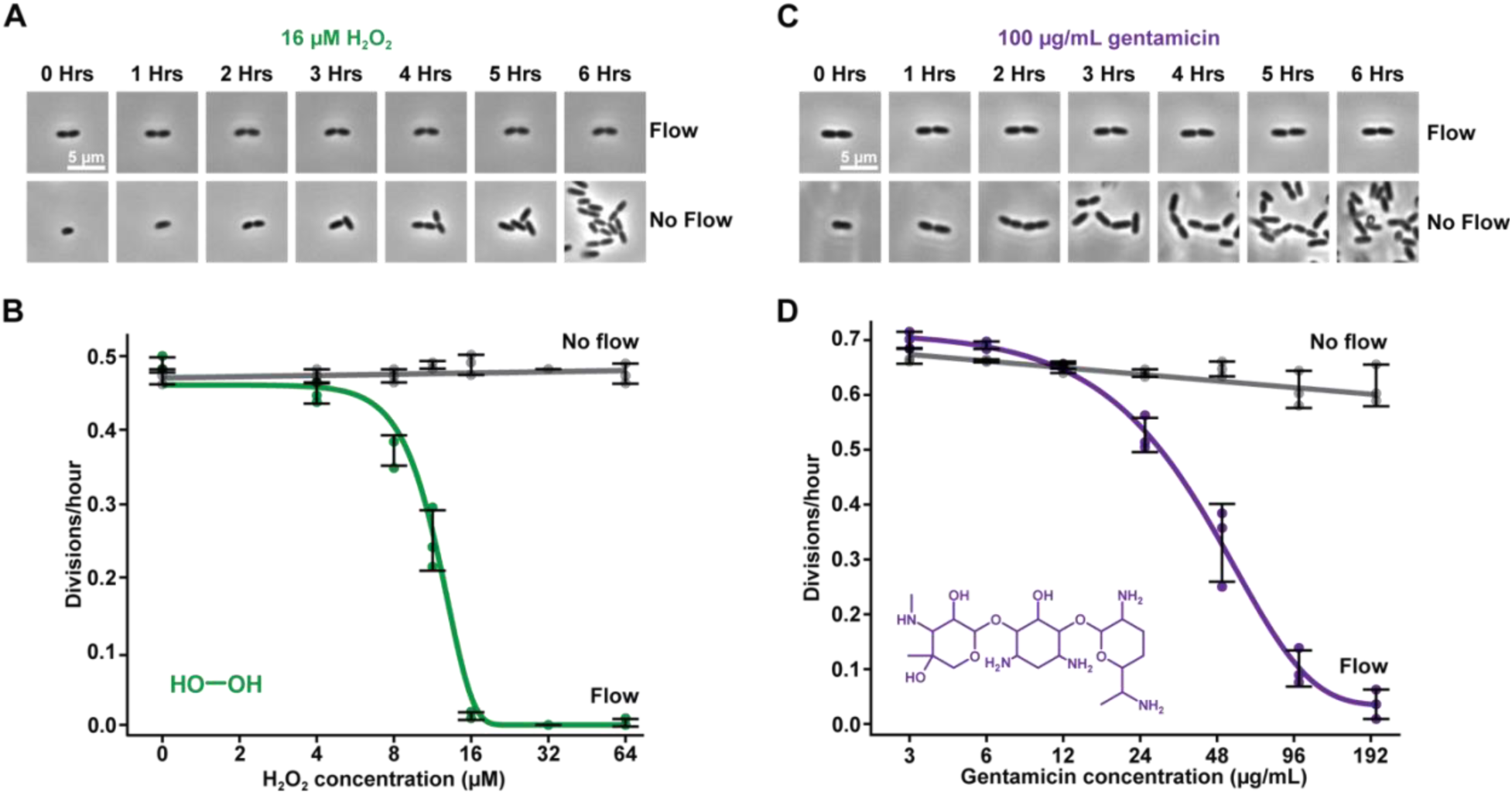
Flow overcomes bacterial resistance to H_2_O_2_ and gentamicin. **(A)** Images of naturally-resistant *P. aeruginosa* cells capable of growth in 16 µM H_2_O_2_ over 6 hours without flow. With flow, cells did not grow with 16 µM H_2_O_2_. **(B)** Growth of *P. aeruginosa* cells in M9 minimal media with flow (green) and without flow (gray) over 2 hours at various H_2_O_2_ concentrations. **(C)** Images of resistant *P. aeruginosa* cells capable of growth in 100 µg/mL gentamicin over 6 hours without flow. With flow, cells did not grow with 100 µg/mL gentamicin. **(D)** Growth of *P. aeruginosa* cells in LB media with flow (purple) and without flow (gray) over 4 hours at various gentamicin concentrations. All growth experiments were performed at a shear rate of 240 sec^-1^ and were imaged near the beginning of a 2 cm channel. For each biological replicate, 28 cells were chosen at random for quantification. Error bars show SD of 3 biological replicates.

Can flow overcome resistance to clinically-relevant antibiotics? Gentamicin is an effective antibiotic towards *P. aeruginosa* that targets protein synthesis^39^. Introduction of a resistance gene encoding an acetyltransferase that directly modifies gentamicin provides *P. aeruginosa* with gentamicin resistance^40^ (Figure S4). To test how flow impacts gentamicin resistance, we treated resistant *P. aeruginosa* cells with varying gentamicin concentrations with and without flow. Without flow, cells were resistant to gentamicin, showing growth at concentrations up to 200 µg/mL (Figure 2C, 2D). With flow, cells became sensitive to gentamicin, as concentrations as low as 25 µg/mL limited growth (Figure 2C, 2D). Together, our results reveal that flow overcomes resistance towards two chemically distinct antimicrobials by replenishing molecules faster than bacteria can neutralize them.

### Physical and chemical properties of antimicrobials tune their delivery

Delivery of drugs to their target microenvironment is often a rate-limiting step in therapeutic intervention^41^. This is particularly evident when small molecules are chemically neutralized during the delivery process, such as by resistant bacteria. One potential solution to this problem is to increase the chemical dose administered. We simulated the impact of increasing H_2_O_2_ dosage across a resistant *P. aeruginosa* population. Our simulations generated the hypothesis that increasing dosage allows deeper delivery into bacterial populations (Figure 3A). Guided by our simulations, we performed growth experiments in 27 centimeter long channels (Figure S1) with increasing H_2_O_2_ doses delivered with a constant flow of 240 sec^-1^ (Figure 3B). As we increased the H_2_O_2_ dose from 8 to 64 µM, we observed that spatial growth gradients continually shifted downstream. These spatial changes in growth closely match our simulations and reveal that increasing dosage promotes delivery deeper into resistant *P. aeruginosa* populations.

**Figure 3:**
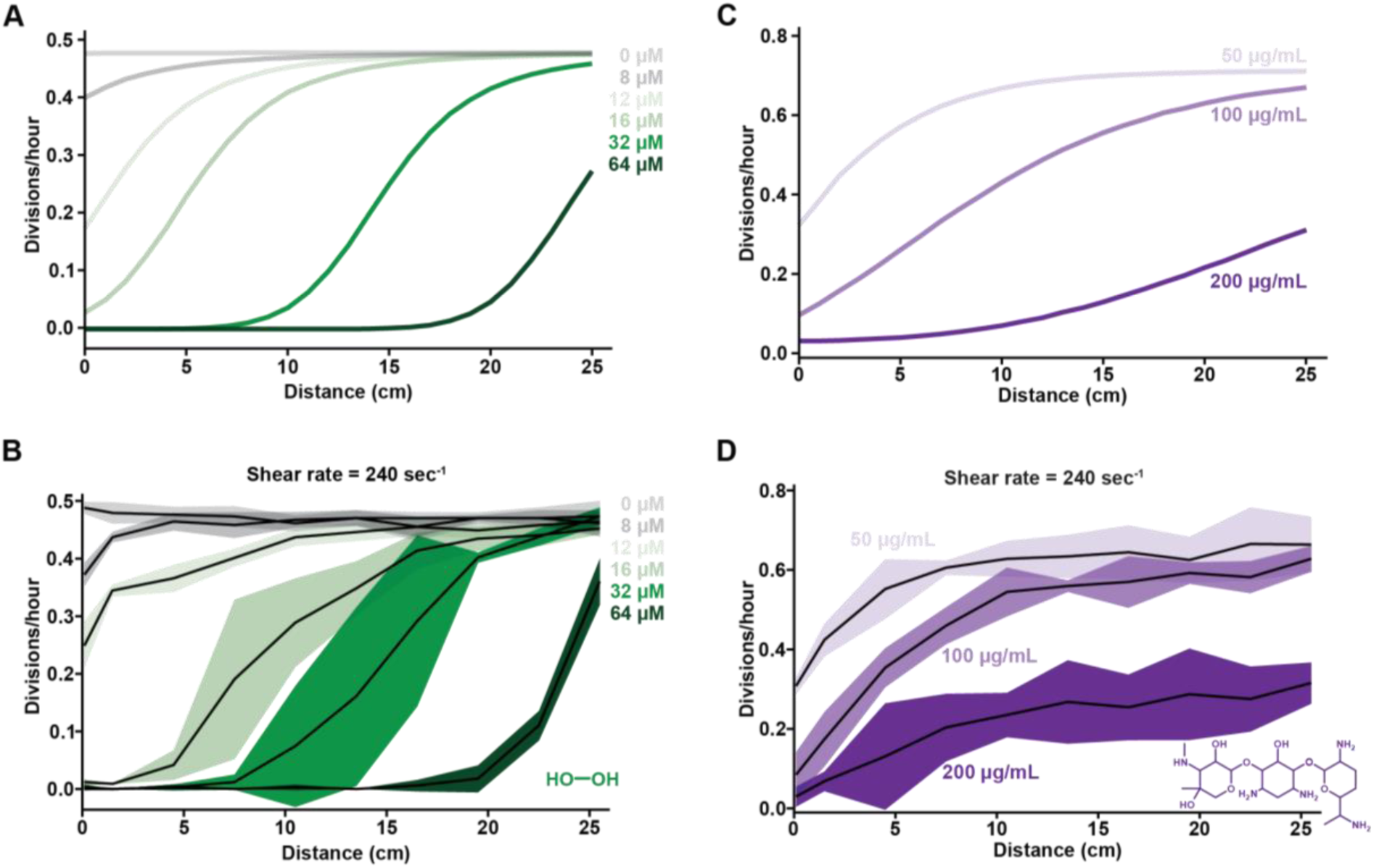
Physical and chemical properties of antimicrobials tune their delivery. **(A)** Simulations of cell growth across a microfluidic channel with increasing concentrations of H_2_O_2_ predict growth gradients shift downstream as concentration increases. **(B)** Experiments measuring *P. aeruginosa* growth closely match the simulations in A. Experiments were performed for 2 hours in M9 minimal media supplemented with various H_2_O_2_ concentrations. **(C)** Simulations of cell growth across a microfluidic channel with increasing concentrations of gentamicin predicts growth gradients shift downstream as concentration increases. The predicted gradients have flatter slopes than those in A for H_2_O_2_, as gentamicin is larger and diffuses more slowly. **(D)** Experiments measuring *P. aeruginosa* growth closely match the simulations in C. Experiments were performed for 4 hours in LB media supplemented with various gentamicin concentrations. All simulations included molecular diffusion, flow, removal, and the relationship between concentration and growth from Figure 2. All growth experiments were performed in 27 cm long channels at a shear rate of 240 sec^-1^. For each biological replicate, 28 cells were chosen at random for quantification. Shaded regions show SD of 3 biological replicates.

How does the physical size of an antimicrobial impact delivery? Antimicrobials come in different sizes, ranging from smaller molecules (such as H_2_O_2_) to larger molecules (such as gentamicin). According to the Stokes-Einstein equation, as molecular size increases, diffusion decreases. Thus, delivery of H_2_O_2_ and gentamicin could differ. To predict if physical size impacts delivery, we simulated the delivery of gentamicin doses (Figure 3C, S6) and compared with our H_2_O_2_ simulations (Figure 3A, S6). Consistent with our hypothesis that diffusion would impact delivery, our gentamicin simulations yielded delivery curves with flatter profiles and lower slopes than our H_2_O_2_ simulations. By simulating molecules with a wide range of diffusion coefficients (Figure S7), we established how molecular size could impact drug delivery into cellular populations. To experimentally test how molecular size impacts delivery, we repeated our long channel growth experiments with constant flow and increasing gentamicin doses (Figure 3D). Supporting our hypothesis, gentamicin generates growth gradients that are flatter than those found for the much smaller antimicrobial H_2_O_2_ (Figure 3D). Collectively, our simulations and experiments demonstrate how chemical dosage and physical size of antimicrobials tune delivery into resistant bacteria populations.

### Flow restores antimicrobial effectiveness across bacterial populations

Antimicrobial resistance often depends on the collective ability of bacterial populations to neutralize molecules. Bacteria can effectively shield one another from antimicrobials and generate spatial growth gradients. Based on our discovery that flow can promote delivery, we reasoned that increasing flow could overcome cell-cell shielding and restore antimicrobial effectiveness throughout the entire population. To test the impact of flow, we treated resistant *P. aeruginosa* populations in 27 centimeter long channels with 16 µM H_2_O_2_ delivered at different shear rates (80, 240, 800 sec^-1^) (Figure 4A). While low flow (80 sec^-1^) did not inhibit any growth, medium flow (240 sec^-1^) generated spatial growth gradients where cells at the start of the channel shielded cells at the end of the channel (Figure 4A). In contrast, high flow (800 sec^-1^) overwhelmed cell-cell shielding and restored the antimicrobial effectiveness of H_2_O_2_. Thus, our results highlight how flow can overcome population-level resistance by enhancing the physical process of delivery.

**Figure 4:**
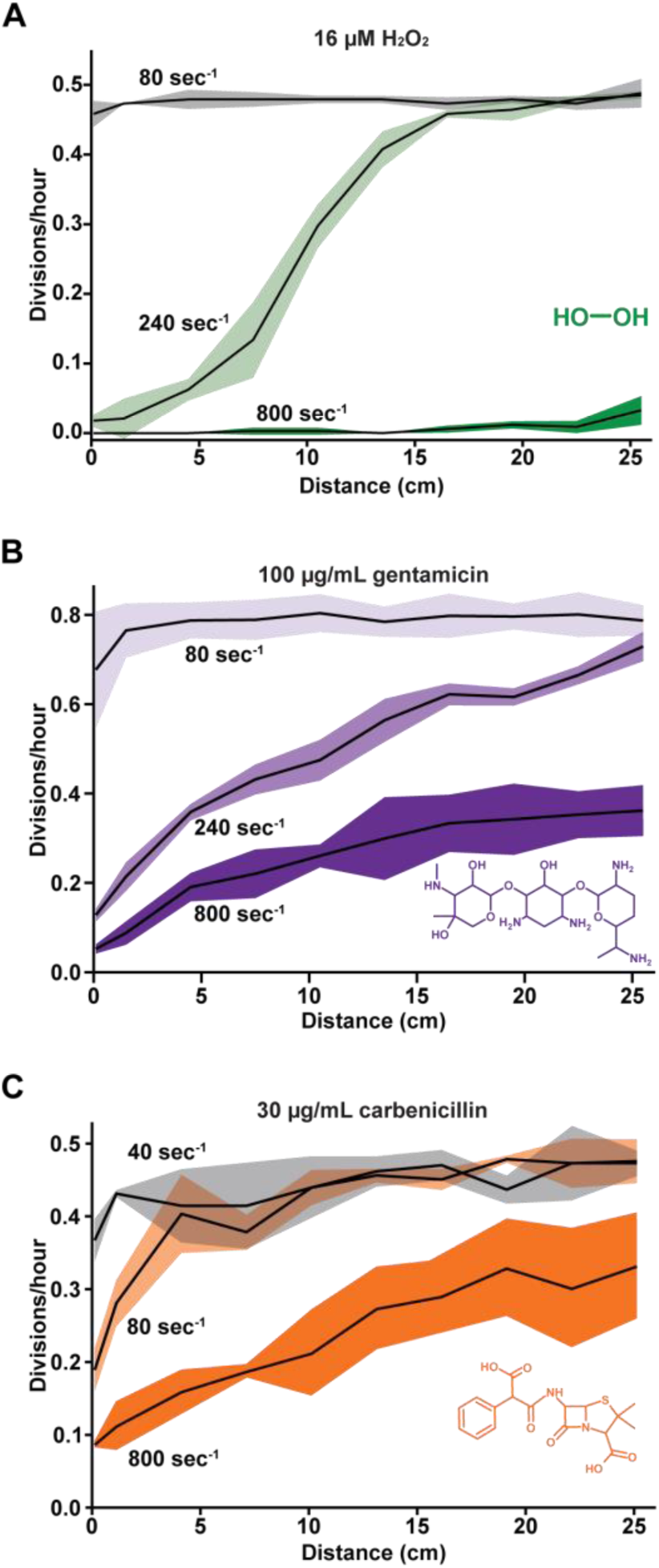
Flow restores antimicrobial effectiveness across bacterial populations. **(A)** Experiments measuring *P. aeruginosa* growth for 2 hours in M9 minimal media with 16 µM H_2_O_2_ at various shear rates. **(B)** Experiments measuring *P. aeruginosa* growth for 4 hours in LB media with 100 µg/mL gentamicin at various shear rates. **(C)** Experiments measuring *P. aeruginosa* growth for 4 hours in LB media with 30 µg/mL carbenicillin at various shear rates. Higher shear rates increase effectiveness of H_2_O_2_, gentamicin, and carbenicillin against resistant *P. aeruginosa* strains. We interpret that growth gradients for gentamicin and carbenicillin have flatter slopes than H_2_O_2_ because H_2_O_2_ is smaller and has a higher diffusion coefficient. All experiments were performed in 27 cm long channels. For each biological replicate, up to 30 cells were chosen at random for quantification. Shaded regions show SD of 3 biological replicates.

To explore if flow can overcome population-level shielding of other antimicrobials, we replicated our experiments with gentamicin. Gentamicin is an aminoglycoside that inhibits protein synthesis^39^, and we used *P. aeruginosa* cells with an acetyltransferase that provided resistance to gentamicin^40^. Using 27 centimeter long channels, we delivered constant doses of 100 µg/mL gentamicin with varied shear rates (80, 240, 800 sec^-1^) (Figure 4B). Similar to H_2_O_2_, low flow (80 sec^-1^) did not inhibit growth, medium flow (240 sec^-1^) generated spatial growth gradients due to cell-cell shielding, and high flow (800 sec^-1^) restored the effectiveness of gentamicin throughout the entire population (Figure 4B). These results reinforce our conclusion that flow can overcome population-level shielding and suggest that this is a broadly conserved biophysical phenomenon.

To further explore the effect of flow on antimicrobial shielding, we extended our experimental approach to the β-lactam antibiotic carbenicillin. Carbenicillin inhibits cell wall biosynthesis^42^, and we used *P. aeruginosa* cells with a β-lactamase that provides resistance to carbenicillin^43^ (Figure S4). Flow increases the sensitivity of cells to carbenicillin (Figure S8), such that 30 µg/mL carbenicillin inhibits cell division and increases cell length (Figure S9). Using 27 centimeter long channels, we delivered constant doses of 30 µg/mL carbenicillin (Figure 4C) with varied shear rates (40, 80, 800 sec^-1^). Similar to H_2_O_2_ and gentamicin, low flow (40 sec^-1^) did not inhibit growth, medium flow (80 sec^-1^) generated spatial growth gradients due to cell-cell shielding, and high flow (800 sec^-1^) restored the effectiveness of carbenicillin throughout the entire population (Figure 4C). Together, our experiments with H_2_O_2_, gentamicin, and carbenicillin demonstrate that flow has a powerful ability to restore antimicrobial effectiveness throughout a resistant bacterial population by boosting delivery.

## Discussion

Our experiments and simulations demonstrate how flow overcomes bacterial resistance. Without flow, bacteria can resist many antimicrobials through degradation or chemical modification. Resistant cells generate local zones of depletion and eventually clear antimicrobials from their environment (Figure 5A). However, as flow increases and replenishes antimicrobials at a rate faster than the cell removal rate, local concentrations increase (Figure 5B). In higher flow, resistant cells do not have time to neutralize their environment, and antimicrobial effectiveness is restored (Figure 5C). Also, flow-enhanced delivery allows antimicrobials to penetrate deeper into target populations, preventing a subset of cells from escaping treatment (Figure 5). Thus, our results reveal a simple biophysical phenomenon that enhances antimicrobial effectiveness and could be leveraged to revitalize antimicrobials that appear neutralized by bacterial resistance.

**Figure 5:**
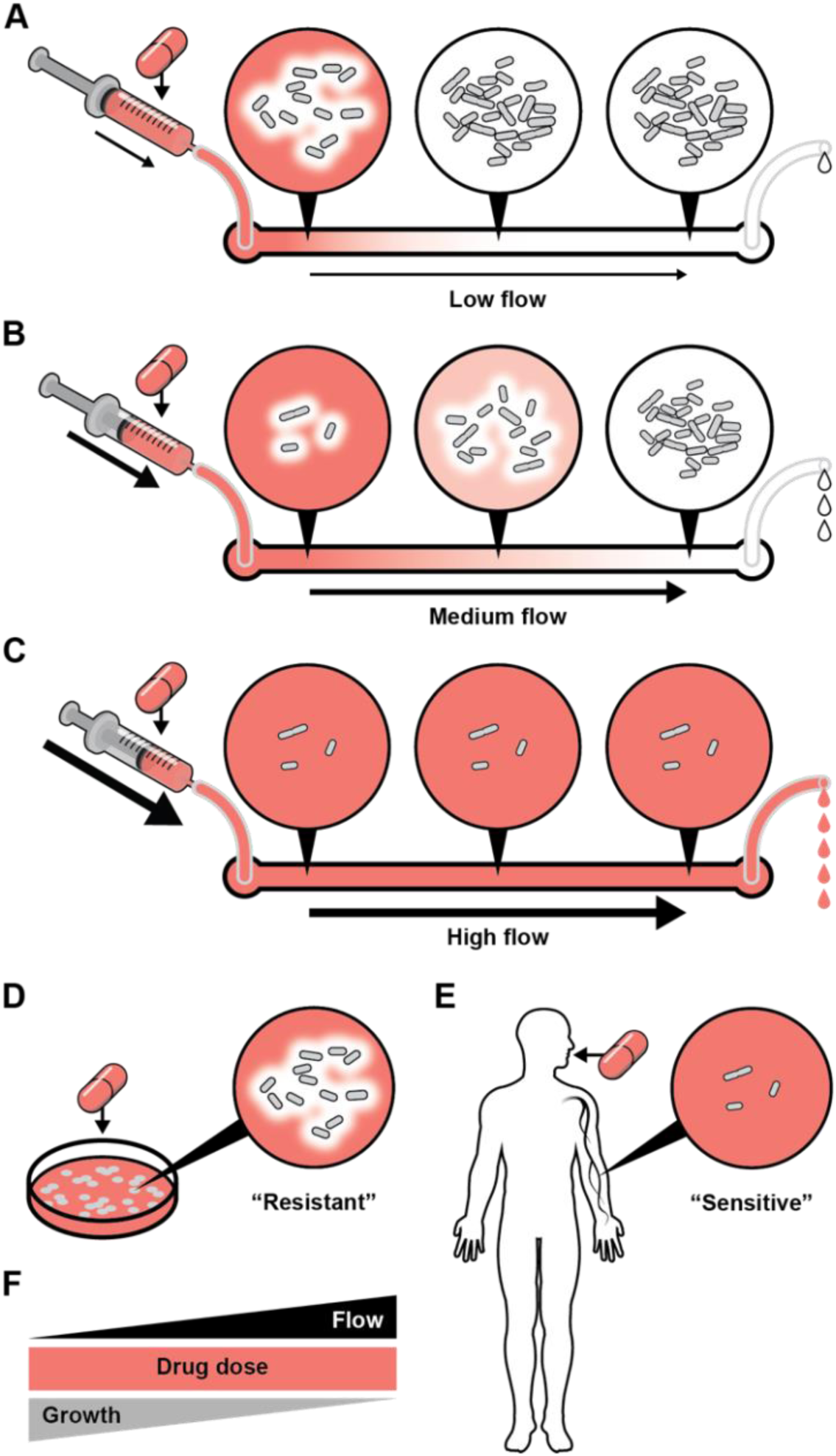
Fluid flow revitalizes antimicrobials by boosting delivery. **(A)** In low flow, cell populations remove antimicrobials faster than delivery, allowing for growth and shielding downstream cells. **(B)** In medium flow, delivery rate is similar to removal rate, allowing antimicrobials to penetrate deeper into populations. Upstream growth is inhibited, but downstream cells can still grow. **(C)** In high flow, delivery rate exceeds removal rate, allowing antimicrobials to penetrate the entire population. Thus, high flow blocks growth of the entire resistant population. **(D)** On petri plates, resistant cells remove antimicrobials, generate zones of clearance, and grow. **(E)** In host-relevant conditions, the same cells experience flow, cannot remove antimicrobials quickly enough, and do not grow. **(F)** When drug dose is held constant, increasing flow prevents cellular growth.

How generalizable is the phenomenon that flow overcomes antimicrobial resistance? As our experiments have been performed with three chemically distinct antimicrobials (H_2_O_2_, gentamicin, carbenicillin), we conclude that the ability of flow to overcome resistance is widely relevant. Importantly, we have shown that flow can enhance effectiveness of two clinically important antibiotic classes: aminoglycosides (represented by gentamicin) and β-lactams (represented by carbenicillin). As gentamicin is modified via a cellular acetylase^40^ and carbenicillin is broken apart by a cellular β-lactamase^42^ (Figure S4), flow can overcome different mechanisms of resistance. Also, flow-enhanced activity does not depend on attributes exclusive to *P. aeruginosa*, indicating that flow should overcome resistance in other bacterial species. Consequently, for any antimicrobial that is chemically destroyed or modified by resistant bacteria, it is logical to infer that flow will increase local concentrations and enhance antimicrobial effectiveness.

Currently, bacterial susceptibility to antibiotics is measured in lab conditions lacking flow. Based on our results, we now realize that the absence of flow in diagnostic assays could significantly misrepresent bacterial susceptibility in host environments, which typically have flow. For example, we have shown the human pathogen *P. aeruginosa* appears resistant to gentamicin in a diagnostic assay without flow, while retaining sensitivity to gentamicin in diagnostic assay with host-relevant flow (Figure 2D). Our results provide proof-of-principle that measuring bacterial resistance in static cultures or petri dishes (Figure 5D) can underestimate antimicrobial effectiveness in host conditions (Figure 5E). Additionally, our work reveals that even when dosage is held constant, drug effectiveness can be significantly enhanced simply by increasing flow (Figure 5F). Thus, we argue that there is a great opportunity to improve bacterial diagnostics through the incorporation of flow and propose that fine-tuning flow could represent a new physical strategy to combat antimicrobial resistance.

## Acknowledgements

We thank Jessica Palalay, Piyush Sharma, Nick Martin, Satish Nair, Jim Imlay, Thomas Kehl-Fie, Wilfred van der Donk, Dan Kearns, and Zemer Gitai for helpful discussions and comments on the manuscript.

## Funding

This work was supported by start-up funds from the University of Illinois at Urbana-Champaign and grant K22AI151263 from the National Institutes of Health to J.E.S.

## Contributions

A.M.S., G.C.P., M.D.K., and J.E.S. designed research. A.M.S., G.C.P., A.S., Z.M., and M.D.K. performed research. A.M.S., G.C.P., A.S., Z.M., M.D.K., and J.E.S. analyzed data. A.M.S. and J.E.S. wrote the paper.

## Supplementary Information

### Materials and Methods

#### Bacterial strains and growth conditions

*Pseudomonas aeruginosa* PA14 and mutant strains used in this paper are described in Supplementary Table 1. *P. aeruginosa* cultures were grown in either LB broth or M9 minimal medium on a cell culture roller drum, and on LB plates (1.5% Bacto Agar) at 37°C. LB broth was prepared using LB broth Miller (BD Biosciences). M9 minimal medium was prepared as 1X M9 salts, 0.4% glucose, 2 mM MgSO_4_, and 100 µM CaCl_2_.

#### Preparation of antimicrobial solutions

LB with H_2_O_2_ was generated as previously described^17^. Briefly, LB was conditioned to remove endogenous H_2_O_2_ and then defined concentrations of H_2_O_2_ were added back. Conditioning was done by diluting *P. aeruginosa* from an overnight culture into LB at a 1:100 ratio, incubating for one hour at 22°C, and filtering to remove cells. H_2_O_2_ solutions were prepared from a 30% w/w stock solution (Sigma-Aldrich). Gentamicin stock solutions were prepared from gentamicin sulfate (VWR) dissolved in water. Carbenicillin stock solutions were prepared from carbenicillin disodium salt (Gold Biotechnology) dissolved in water.

#### Fabrication of microfluidic devices

Microfluidic devices were prepared as previously described^17^. Briefly, photomasks were designed on Illustrator (Adobe Creative Suite) and printed by Artnet Pro, Inc. Using soft lithography techniques, photomask patterns were transfered onto 100 mm silicon wafers (University Wafer) that were spin coated with SU-8 3050 photoresist (MicroChem). Microfluidic devices are made of Polydimethylsiloxane (Dow SYLGARD 184 Kit) at a 1:10 ratio and plasma-treated to bond on a 60 mm x 35 mm x 0.16 mm superslip micro cover glass (Ted Pella, Inc.). Growth experiments used channels with dimensions of 500 µm wide x 50 µm tall x 2 cm long or 500 µm wide x 50 µm tall x 27 cm long with 8 turns. *fro* reporter experiments used channels with dimensions of 500 µm wide x 50 µm tall x 100 cm long with 32 turns.

#### Phase contrast and fluorescence microscopy

Timelapse images were captured on a Nikon ECLIPSE Ti2-E inverted microscope using the NIS Elements interface. For all imaging, we used a Nikon 40x Plan Ph2 0.95 NA objective, a Hamamatsu ORCA-Flash 4.0 LT3 Digital CMOS camera, and a Lumencor SOLA Light Engine LED light source.

#### Preparation of microfluidic devices with bacteria

Microfluidic devices were loaded with bacteria as previously described^17^. Briefly, all experiments were performed at approximately 22°C and with mid-log bacterial cultures. Cultures were loaded into microfluidic devices via syringe (BD Biosciences) on a syringe pump (KD Scientific Legato 210) set at 10 µL/min. Cells were allowed to adhere for 10 minutes before experimental flow exposure. Device inlets are attached to a length of polyethylene tubing, ID 0.015” x OD 0.043” (Brain Tree Scientific) that is sheathed over a 26-gauge x 1/2” hypodermic needle (Air-tite Products) that is affixed on a syringe. Device outlets contain another length of tubing for vacating spend media into a bleach-containing waste container. Media-filled syringes were fastened on a syringe pump that was set at flow rates between 0.5 - 10 µl/min, which correspond with shear rates between 40 - 800 sec^-1^.

#### Shear rate calculations

Shear rate experienced in microfluidic devices was calculated using this equation:

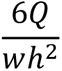

Where *Q* is flow rate, *w* is channel width, and *h* is channel height. Shear rate is specified throughout in units of sec^-1^.

#### Quantification of *fro* expression

Expression of *fro* reporter activity was quantified as previously described^17^. Briefly, images were captured every 30 minutes for 4 hours. Quantification used a MATLAB (Mathworks)-based program to identify and quantify single cell fluorescence intensity (OUFTI). YFP fluorescence was used to quantify *fro* expression and mCherry fluorescence was used as a normalization control. Reporter strains expressed mCherry from a chromosomally encoded constitutive promoter (*P_A1/04/_*_03_)^18^. *fro* reporter activity was quantified as the ratio of YFP to mCherry. Experiments measuring *fro* expression used wildtype reporter cells (JS16) or Δ*pilA* mutant reporter cells (JS27).

#### Quantification of cellular growth

To measure cell growth during antimicrobial treatments with and without flow, images were captured every 5 minutes for up to 6 hours. Cell growth quantification was performed manually on ImageJ software by counting total divisions and dividing by time. At least 28 cells per field were chosen at random for growth tracking. All growth experiments were performed with cells lacking *pilA*, which prevented twitching motility and allowed for easy cell tracking. H_2_O_2_ and carbenicillin growth experiments used strain JS176, which contained an unmarked Δ*pilA*::FRT mutation. Gentamicin growth experiments used strain JS177, which contained a Δ*pilA*::*aacC1* mutation that provided resistance to gentamicin. H_2_O_2_ growth experiments were performed in M9 media with glucose. Gentamicin and carbenicillin growth experiments were performed in LB media.

#### Biophysical simulations

Simulations of molecular advection and diffusion were performed as previously described^17^. Briefly, simulated channels were randomly filled with molecules that were assigned diffusive behavior by combining laminar transport and Brownian dynamic simulations. Channel parabolic flow speed profiles were modeled using the Hagen-Poiseuille equation. Molecules were allowed to diffuse along the x-axis and y-axis of the channel. H_2_O_2_ was simulated with a diffusion coefficient of 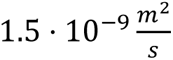. Molecular removal was estimated to be 1 of every 100 molecules that reached the channel bottom (where cells are located). Gentamicin simulations were performed using flow and removal parameters matching the H_2_O_2_ simulations. The diffusion coefficient used for gentamicin was 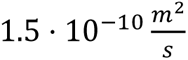, which is 10-fold less than H_2_ O_2_, accounting for the molar mass of gentamicin being approximately 10-fold greater than H_2_O_2_. Growth simulations were generated by combining concentration simulations with the quantitative relationships between concentration and growth established in Figure 2.

---

**Figure S1:**
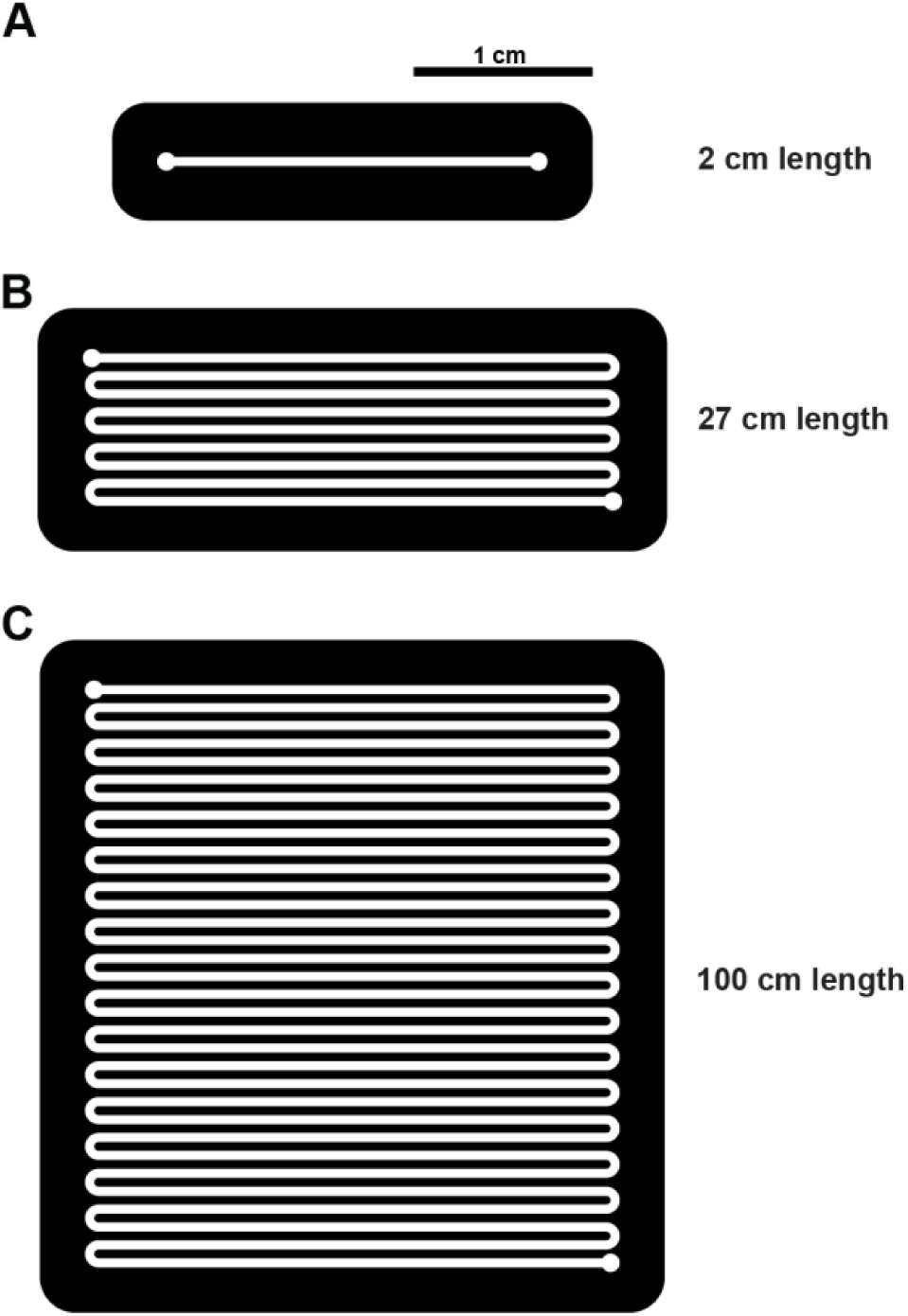
Designs of microfluidic devices used in this study. **(A)** Top-down view of a 2 cm long device. **(B)** Top-down view of a 27 cm long device containing 8 turns. **(C)** Top-down view of a 100 cm long device containing 32 turns. Scale bar is 1 cm.

**Figure S2:**
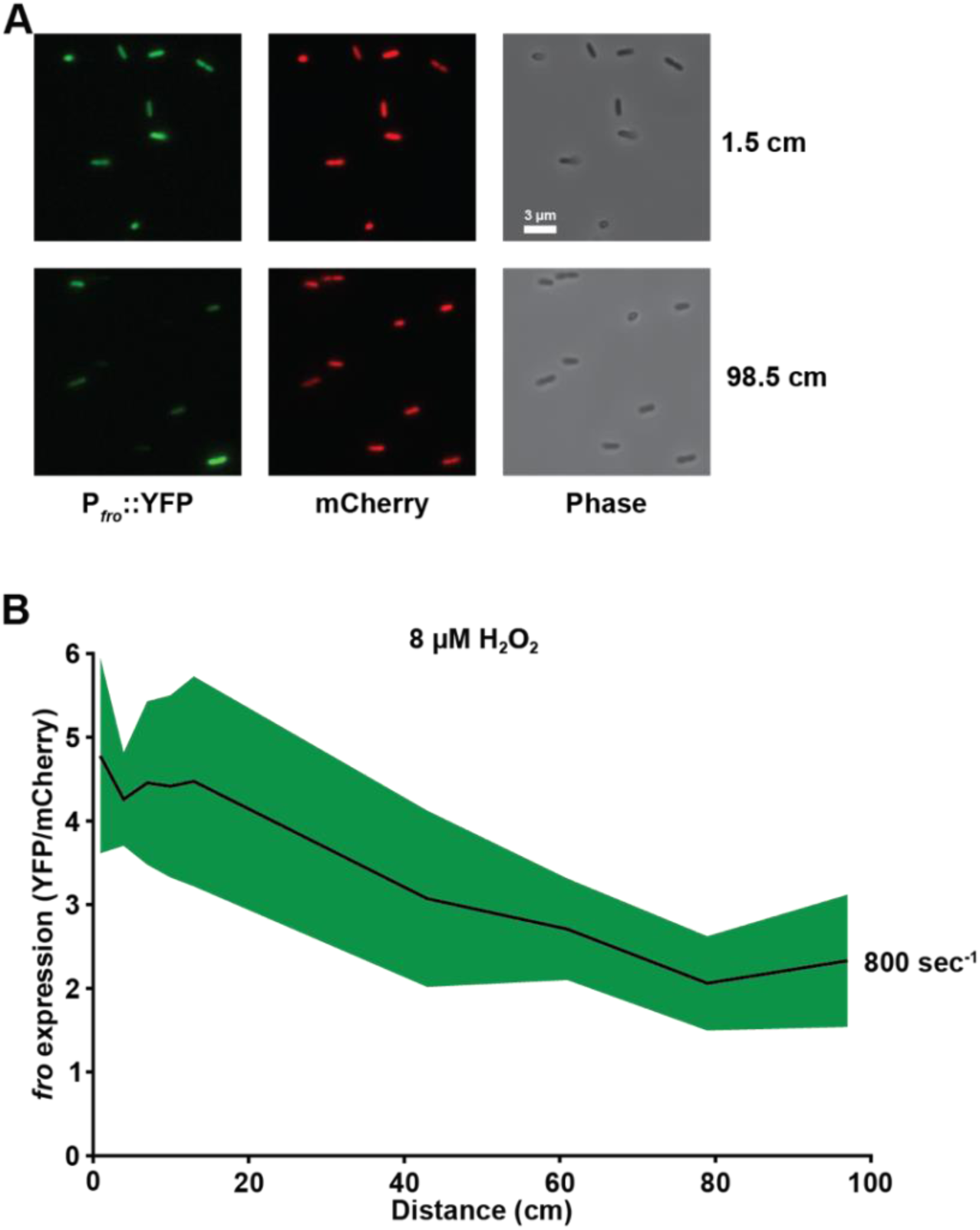
*P. aeruginosa* spatial *fro* gene expression from wildtype cells. **(A)** YFP fluorescence, mCherry fluorescence, and phase contrast images representative of 3 biological replicates of *P. aeruginosa* cells near the beginning (1.5 cm) and end (98.5 cm) of a 100 cm long microfluidic channel. Scale bar is 3 µm**. (B)** *P. aeruginosa fro* reporter expression (which serves as a H_2_O_2_ biosensor) after 3 hours at a shear rate of 800 sec^-1^ with LB media containing 8 µM H_2_O_2_. *fro* expression is quantified as the ratio of YFP to mCherry. Shaded regions show SD of three biological replicates.

**Figure S3:**
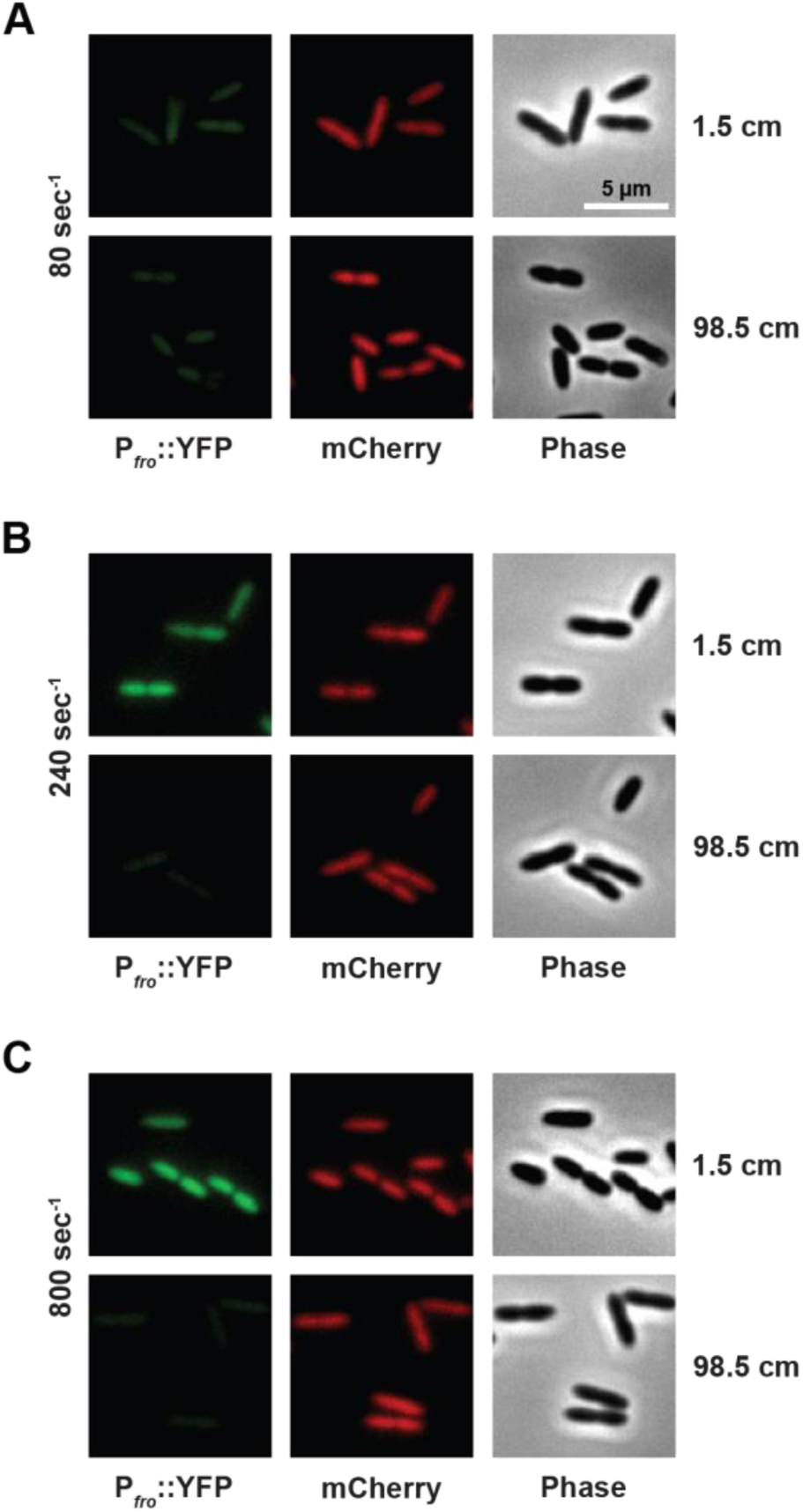
Representative images of H_2_O_2_ spatial gradients. YFP fluorescence, mCherry fluorescence, and phase contrast images representative of three biological replicates of *P. aeruginosa* Δ*pilA* reporter cells at various shear rates of 80 sec^-1^ **(A)**, 240 sec^-1^ **(B)**, and 800 sec^-1^ **(C)**. Images were taken near the beginning (1.5 cm) and end (98.5 cm) of a 100 cm long microfluidic channel. YFP signal decreases over distance as H_2_O_2_ is removed by cells. Cells are exposed to flow with conditioned LB media supplemented with 8 µM H_2_O_2_. Scale bar is 5 µm.

**Figure S4:**
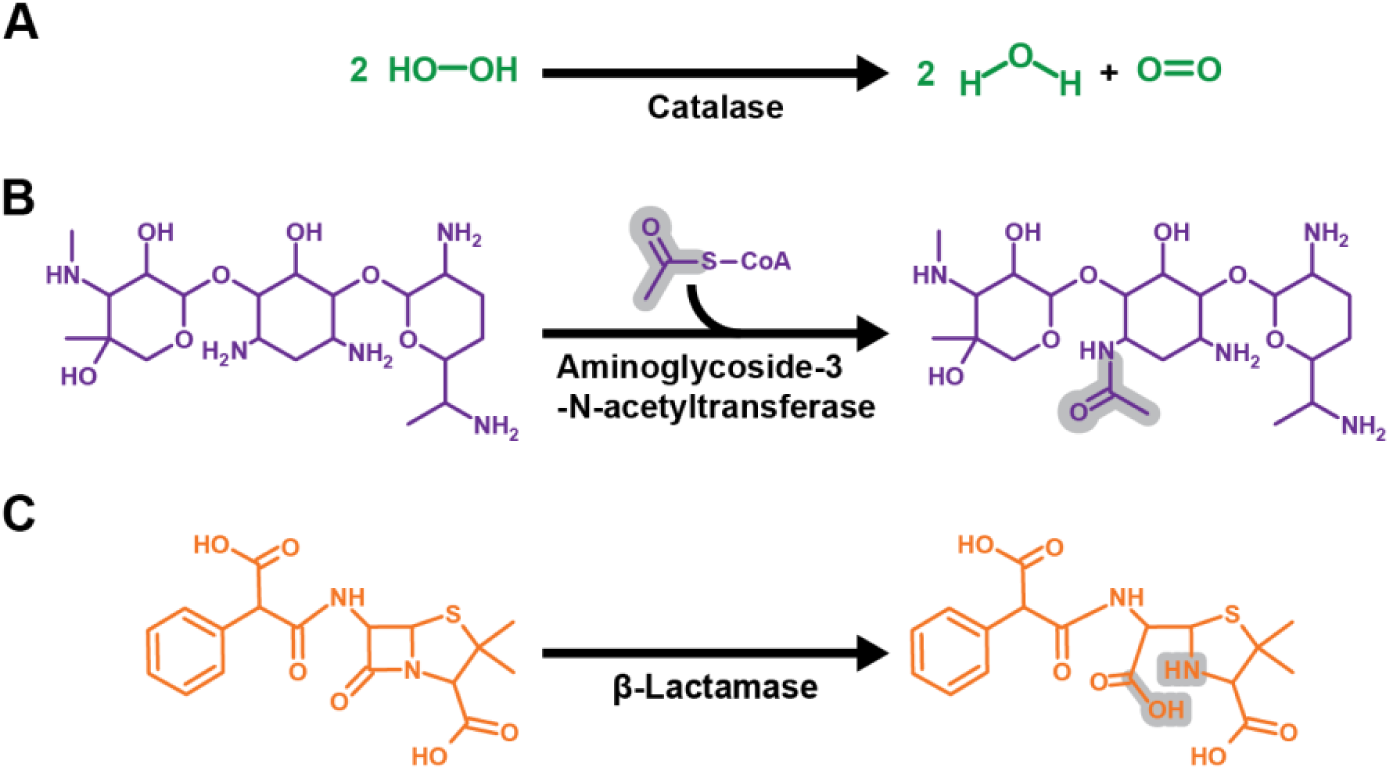
Antimicrobial resistance mechanisms in this study. **(A)** H_2_O_2_ is degraded by catalase into water and oxygen^38^. **(B)** Gentamicin is modified by aminoglycoside-3-N-acetyltransferase that transfers the acetyl group from acetyl-CoA to the 3C amino group of gentamicin^40^. **(C)** Carbenicillin is deactivated by β-lactamase that breaks the amide bond within the β-lactam ring^42^.

**Figure S5:**
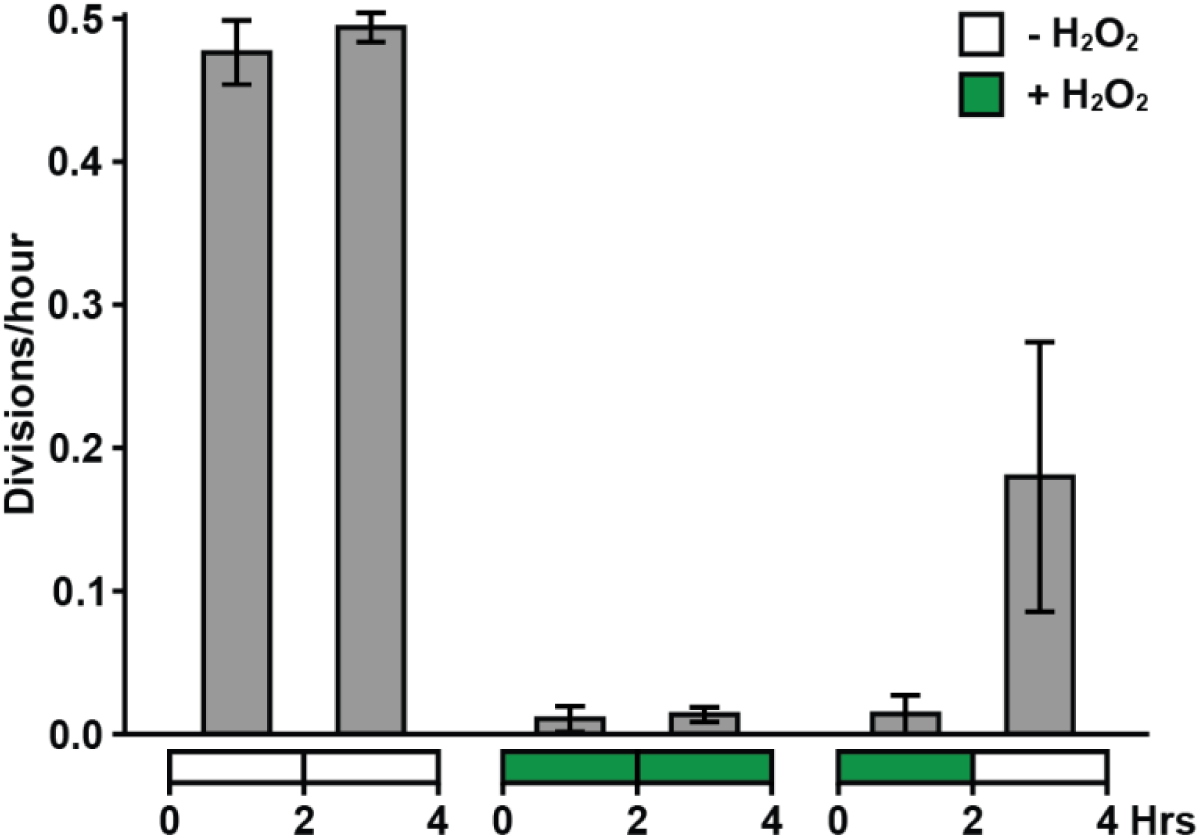
H_2_O_2_ acts as a bacteriostatic antimicrobial towards *P. aeruginosa*. *P. aeruginosa* cell growth in M9 minimal media without H_2_O_2_ (white) or with 16 µM H_2_O_2_ (green) over 4 hours in 240 sec^-1^. Cells that experience no H_2_O_2_ for 4 hours grow normally, and cells exposed to H_2_O_2_ for 4 hours experience growth inhibition. Cells that experience 16 µM H_2_O_2_ for 2 hours before being switched to no H_2_O_2_ for an additional 2 hours exhibit no growth followed by growth. Growth is imaged in a 2 cm channel. For each biological replicate, 28 cells were chosen at random for quantification. Error bars show SD of 3 biological replicates.

**Figure S6:**
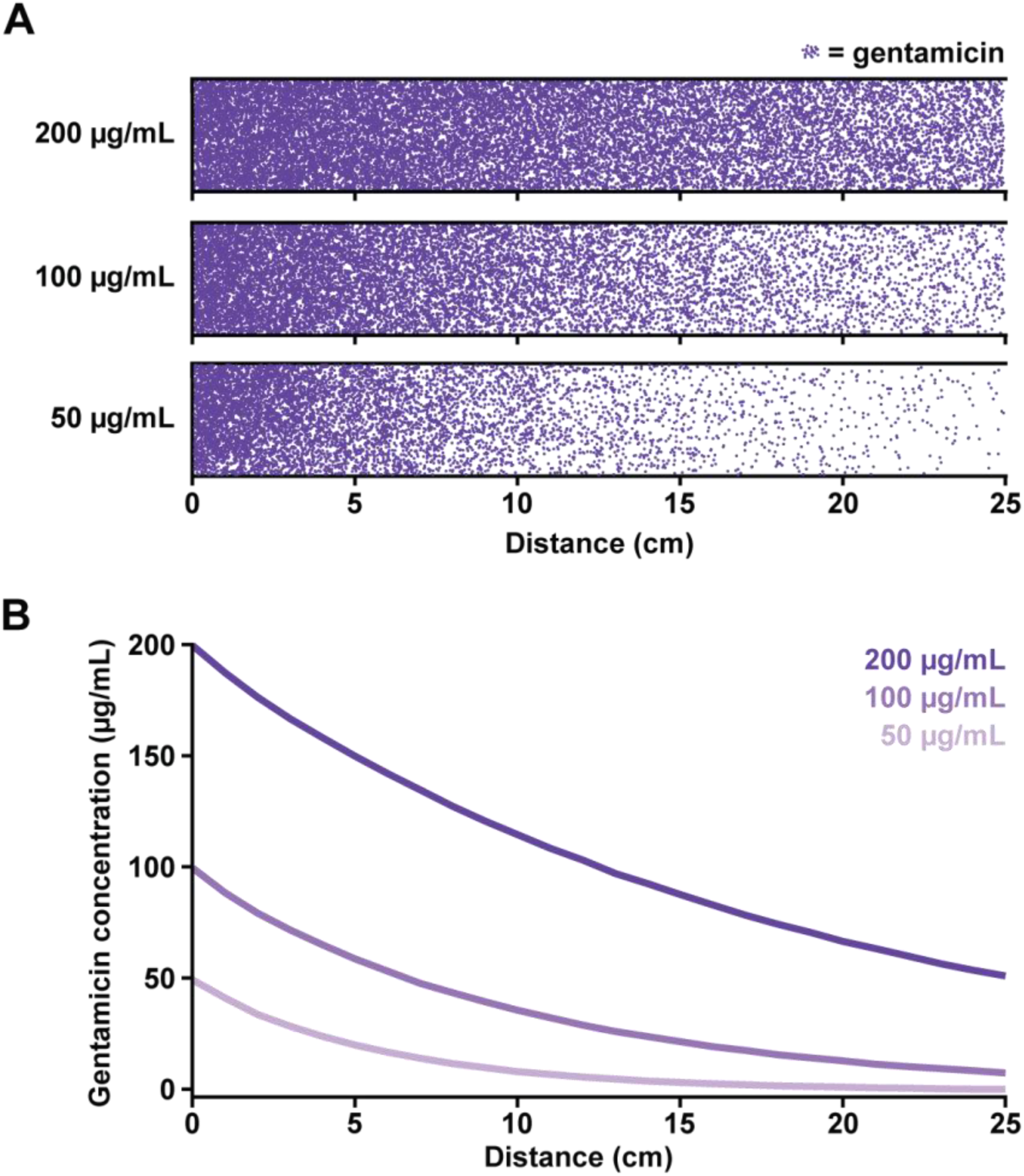
Biophysical simulations predict spatial gentamicin gradients. **(A)** Simulation of gentamicin concentration with respect to distance at various concentrations. Purple dots represent individual gentamicin molecules that experienced diffusion, flow from left to right, and removal when they hit the bottom of the channel to represent bacterial removal. **(B)** Quantification of gentamicin concentration across the length of simulated channels.

**Figure S7:**
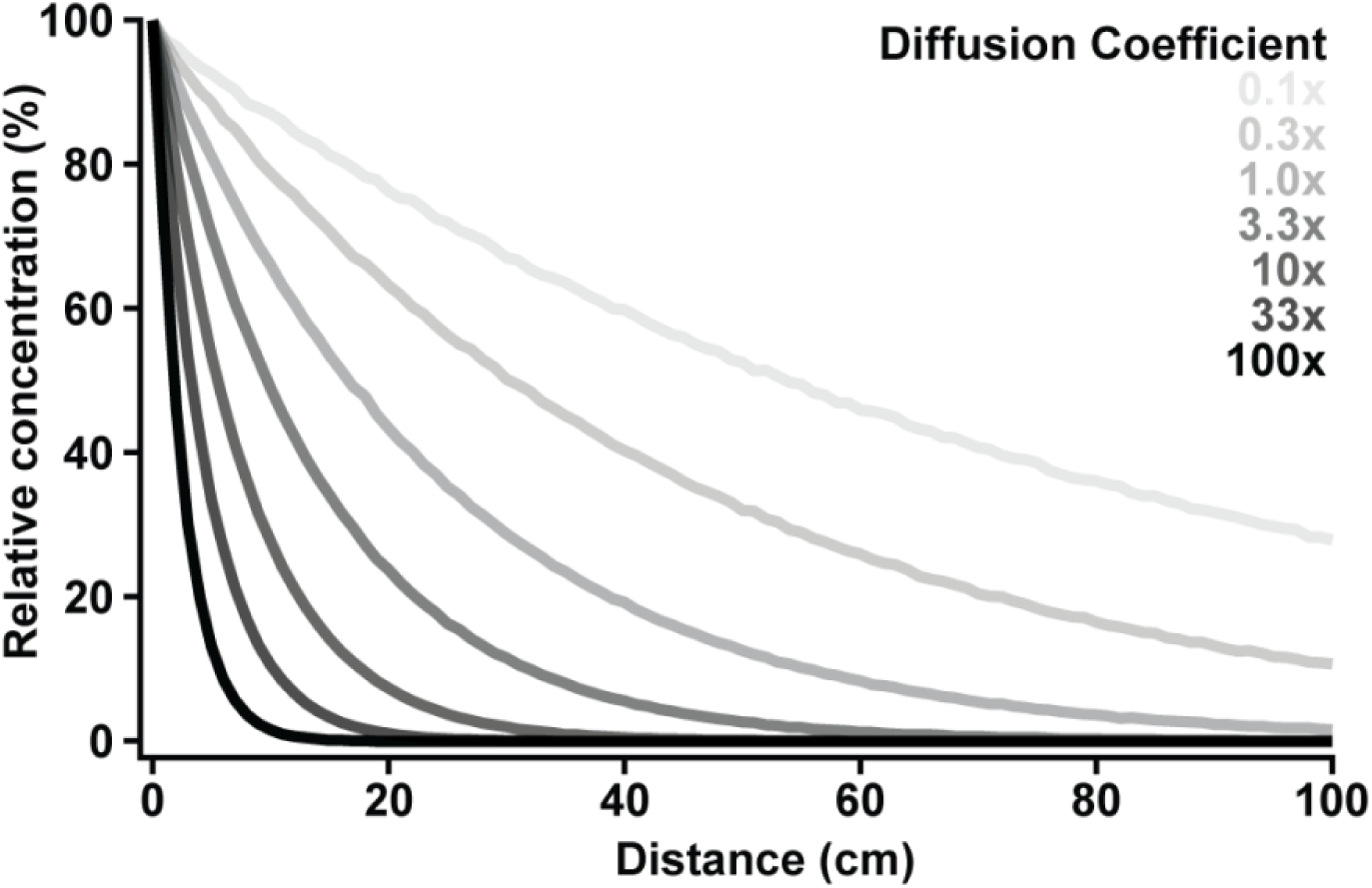
Increasing diffusion generates steeper spatial gradients. Biophysical simulations of molecules with different diffusion coefficients. By holding flow and removal constant, we directly tested how changing diffusion in our simulations would impact the shape of spatial concentration gradients. Compared to 1.0x diffusion (which represents H_2_O_2_), lower diffusion coefficients yield flatter slopes and higher diffusion coefficients yield steeper slopes.

**Figure S8:**
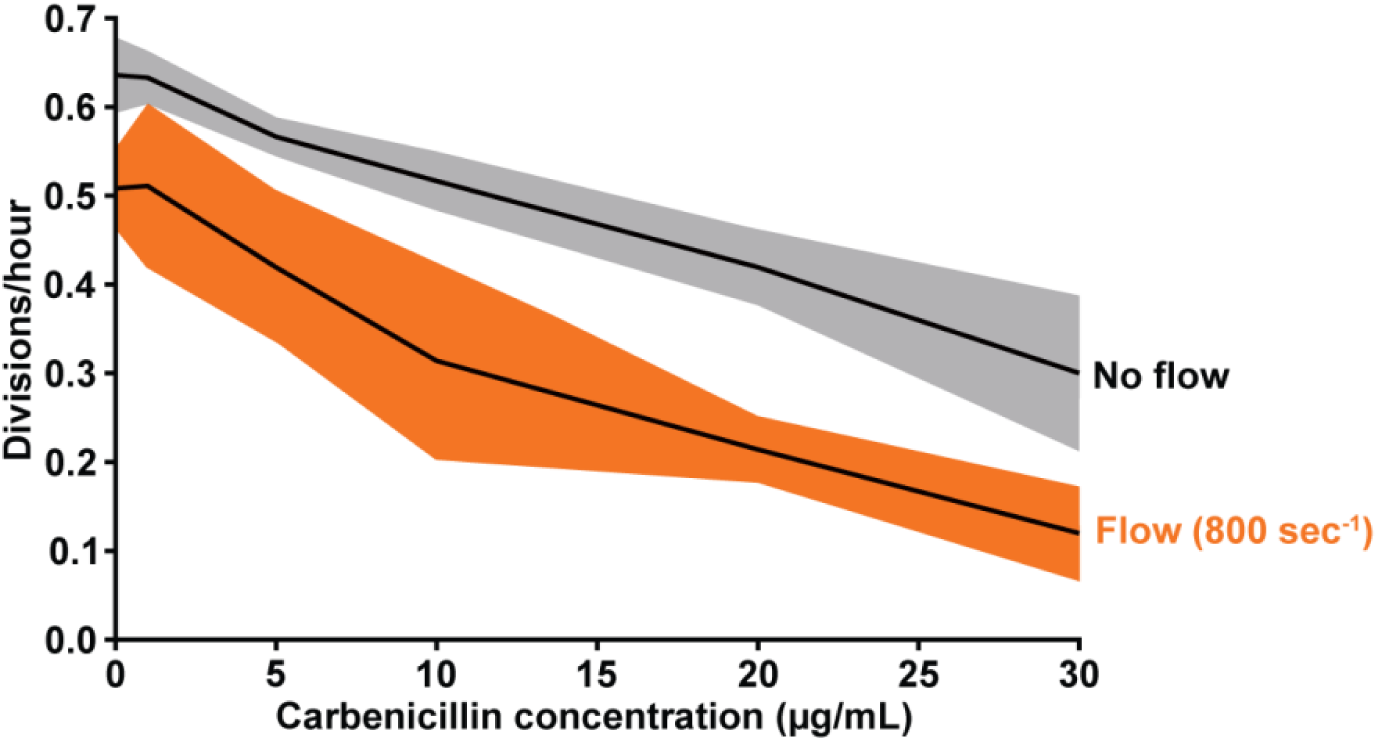
Flow enhances sensitivity of *P. aeruginosa* to carbenicillin. *P. aeruginosa* cell growth over 4 hours in LB media supplemented with various concentrations of carbenicillin. Growth measurements with 800 sec^-1^ flow (orange) and without flow (grey) show cell growth is reduced in flow across all concentrations. For each biological replicate, 30 cells were chosen at random for quantification. Shaded regions show SD of 3 biological replicates.

**Figure S9:**
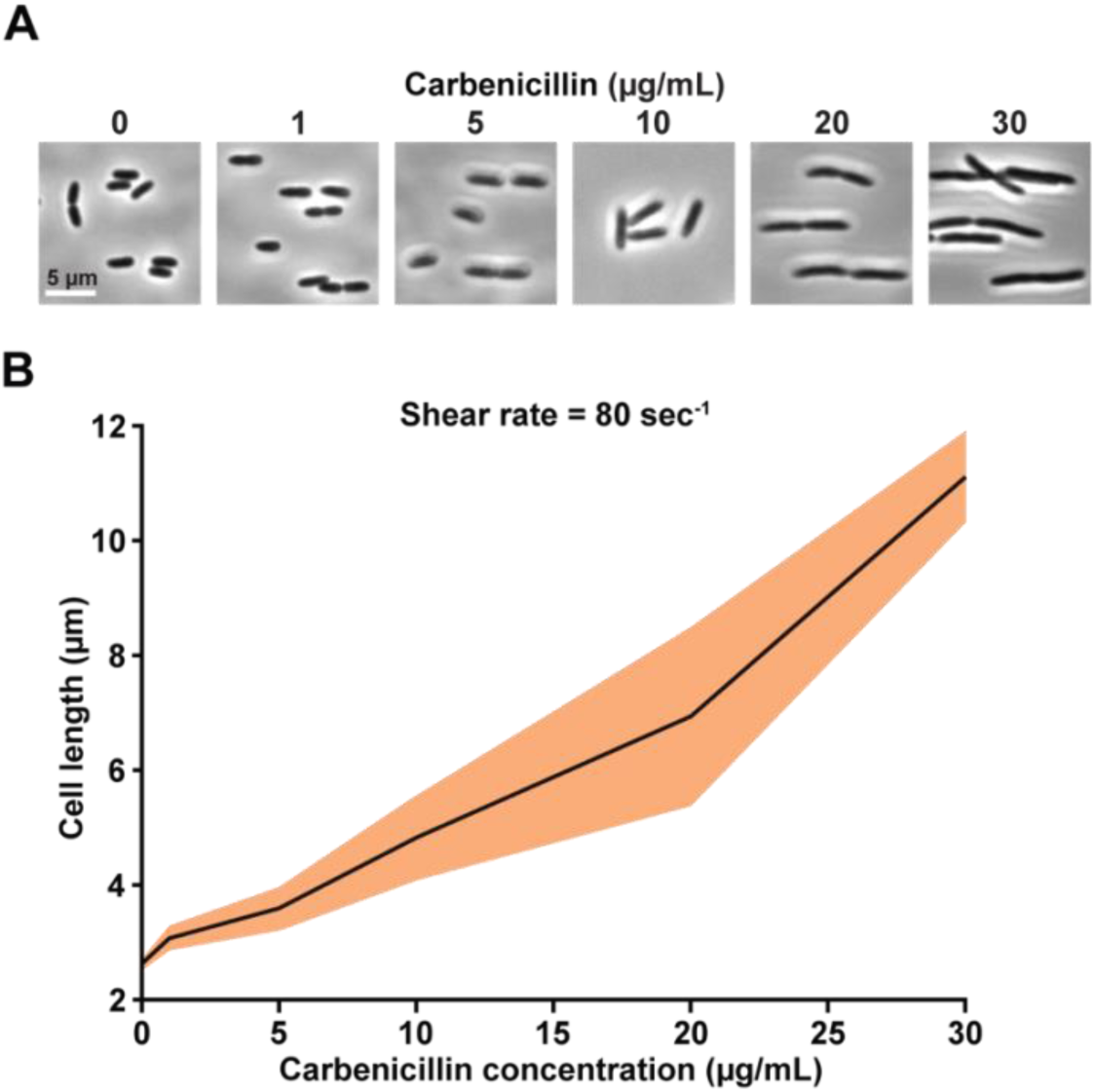
*P. aeruginosa* cells are longer when treated with carbenicillin. **(A)** *P. aeruginosa* cells exposed to increasing concentrations of carbenicillin are longer. Scale bar is 5 µm. **(B)** Quantification of the relationship between *P. aeruginosa* cell length and carbenicillin concentration. For each biological replicate, 30 cells were chosen at random for quantification. Shaded regions show SD of 3 biological replicates.

**Table S1:** *P. aeruginosa* strains used in this study.

| *P. aeruginosa*<br>strains | Description | Source |
| --- | --- | --- |
| PA14 | Wildtype; clinical isolate from burn wound | 44 |
| JS16 | PA14 *froA*::*YFP FRT attB*::[*P*<sub>A1/04/03</sub>-*mCherry FRT*] | 18 |
| JS27 | PA14 *froA*::*YFP FRT attB*::[*P*<sub>A1/04/03</sub>-*mCherry FRT*]Δ*pilA*::*FRT* | 18 |
| JS176 | PA14 Δ*pilA*::*FRT* | 10 |
| JS177 | PA14 Δ*pilA*::*aacC1* | 10 |

## Notes

### Competing Interest Statement

The authors have declared no competing interest.

